# Mosquito odorant receptor sensitive to natural spatial repellents and inhibitory compounds

**DOI:** 10.1101/2022.01.19.476929

**Authors:** Pingxi Xu, Young-Moo Choo, Sunny An, Gabriel M. Leal, Walter S. Leal

**Affiliations:** Department of Molecular and Cellular Biology, University of California-Davis Davis, CA 95616, U.S.A.

**Keywords:** Culex quinquefasciatus, CquiOR27, CquiOR32, γ-octalactone, γ-decalactone, γ-dodecalactone

## Abstract

Previously, we have identified an odorant receptor (OR) from the southern house mosquito *Culex quinquefasciatus*, CquiOR32, which responded to both odorants (agonists) and inhibitory compounds (antagonists). CquiOR32/CquiOrco-expressing oocytes responded to methyl salicylate and other odorants with inward (regular) currents but gave currents in the reverse direction when challenged with eucalyptol and other inhibitors. To determine whether hitherto unknown ORs show this intrareceptor inhibition, we have now examined two other receptors in the same cluster, CquiOR27 and CquiOR28. We cloned and tested four variants of CquiOR28, but none of the 250 compounds in our panel of odorants, including an Orco ligand candidate (OCL12), elicited inward or outward currents. By contrast, CquiOR27/CquiOrco-expressing oocytes gave robust, dose-dependent inward currents when challenged with γ-octalactone and other odorants. On the other hand, octylamine and other phenolic compounds elicited dose-dependent currents in the reverse direction. When stimulatory and inhibitory compounds were presented in binary mixtures, γ-octalactone-elicited inward currents were attenuated in a dose-dependent manner according to the concentration of octylamine. As part of our chemical ecology approach, we tested the repellency activity of the best ligands in the surface landing and feeding assay and a newly reported hand-in cage assay. Protection elicited by γ-octalactone did not differ significantly from that of DEET at the same dose. In the hand-in cage assay, a cream formulation of γ-octalactone showed 97.0 ± 1.3% protection, with 47.6 ± 8.3% and 1.4 ± 0.7% landings per trial in the hands covered with a control and γ-octalactone cream, respectively (N=8, p=0.0078, Wilcoxon matched-pairs signed-rank test).

## 1. Introduction

Integration of chemical signals at the insect’s peripheral olfactory system is one of the least understood mechanisms of insect olfaction. There is growing evidence in the literature that in moths (Kaissling, 2014), beetles (Nikonov and Leal, 2002), vinegar fly (Su et al., 2012), mosquitoes (Tauxe et al., 2013), and other insect species, the firing of one odorant receptor neuron (ORN; also referred to as olfactory sensory neuron, OSN), interferes with the signaling of other ORNs. There is convincing evidence that in the vinegar fly, lateral inhibition of collocated ORNs may be mediated by ephaptic coupling (Su et al., 2012). Still, other mechanisms seem to play a role in signal integration. For example, inhibition may occur within the same neuron with the same compound eliciting excitatory and inhibitory responses (de Brito Sanchez and Kaissling, 2005). Recently, we discovered an odorant receptor from the Southern house mosquito, *Culex quinquefasciatus*, CquiOR32, which responded to some compounds with regular inward currents, whereas other compounds elicited currents in the reverse direction (outward currents) (Xu et al., 2019). Specifically, CquiOR32/CquiOrco-expressing oocytes responded in a dose-dependent manner to methyl salicylate and other odorants with inward currents, whereas inhibitory compounds elicited currents in the reverse direction (outward currents). When CquiOR32/CquiOrco-expressing oocytes were challenged with mixtures of stimulatory and inhibitory compounds, inward currents were attenuated in a dose-dependent fashion. The higher the inhibitor dose, the smaller the inward currents; binary mixtures with high concentrations of inhibitors elicited currents in the reverse direction. The data thus suggest that intrareceptor inhibition occurs, although it remains to be determined whether odorant (stimulant) and inhibitor bind to the same or different binding sites in CquiOR32 (Xu et al., 2019). Additionally, we identified an orthologue of CquiOR32 in the genome of the yellow fever mosquito, *Aedes aegypti*, AaegOR71, which showed intrareceptor inhibition, although no inhibitor-elicited outward currents could be recorded (Xu et al., 2019).

Phylogenetic analysis suggests that three other *Cx. quinquefasciatus* ORs belong to a CquiOR32 cluster: CquiOR36, CquiOR27, and CquiOR28 (Taparia et al., 2017). Previously, we have deorphanized CquiOR36, which is narrowly tuned to acetaldehyde and showed no evidence of intrareceptor inhibition (Choo et al., 2018). We have now cloned CquiOR27 and CquiOR28 and attempted to deorphanize these receptors using the *Xenopus* oocyte recording system. CquiOR28/CquiOrco-expressing oocytes did not respond to any of the 250 compounds in our panel of odorants, including the Orco ligand candidate OCL12 (Chen and Luetje, 2012) (also known as VUAA3 (Taylor et al., 2012)). On the other hand, we recorded robust inward currents from CquiOR27/CquiOrco-expressing oocytes challenged with γ-octalactone and other odorants. By contrast, octylamine and other phenolic compounds elicited currents in the reverse direction and, when presented as binary mixtures, attenuated the responses to γ-octalactone in a dose-dependent manner. As part of our reverse chemical ecology approach (Leal, 2005), we measured the behavioral responses of *Cx. quinquefasciatus* to γ-octalactone in repellency assays. Protection data obtained with the surface landing and feeding assay (Leal et al., 2017; Xu et al., 2014) and a newly reported hand-in cage assay suggest that γ-octalactone is a potent mosquito repellent.

## 2. Material and methods

### 2.1. Insect preparations

*Cx. quinquefasciatus* used in this study were from a laboratory colony started in the 1950s from adult mosquitoes collected in Merced, CA. This colony has been maintained at the Kearney Agricultural Research Center, University of California, Parlier, CA, and kept for almost ten years in Davis. Mosquitoes were maintained at 27 ± 1°C, 75 ± 5% relative humidity, and under a photoperiod of 12:12 h.

### 2.2. RNA and DNA extraction, DNA synthesis, and cloning

Total RNA was extracted from one thousand 4-7 day-old female *Cx. quinquefasciatus* antennae with TRIzol reagent (Invitrogen, Carlsbad, CA). Antennal cDNA was synthesized from 1 μg of antennal total RNA using SMARTer™ RACE cDNA amplification kit according to manufacturer’s instructions (Clontech, Mountain View, CA). To obtain full-length coding sequences, PCRs were performed using the following gene-specific primers:

CqOR27-Fw: 5’-ATGGACACAGTCCGGTGGGCTTCA-3’;

CqOR27-Rv: 5’-TTACAACTTGTTAAGCAAATCTTTGAGAACCAAATAGTACGAG -3’

CqOR28-Fw: 5’-ATGGACGCCACCCAGCGAATCAAA-3’;

CqOR28-Rv: 5’-TTACAGTTTGCTCAAAACGTCATTCAACACCAAATA-3’.

PCR products were purified by a QIAquick gel extraction kit (Qiagen, Germantown, MD) and then cloned into pGEM-T vector (Promega, Madison, WI). Plasmids were extracted by a QIAprep spin miniprep kit (Qiagen) and sequenced. CquiOR27 and CquiOR28 were subcloned separately from pGEM-T-CquiOR27 and pGEM-T-CquiOR28 into pGEMHE using In-Fusion HD Cloning Kit according to the manufacturer’s instructions (Clontech). PCR In-Fusion primers were designed according to the user manual. CquiOR27-In-Fusion-Fw primer:

5’-AGATCAATTCCCCGGG<u>ACC</u>ATGGACACAGTCCGGTGG-3’ and

CquiOR27-In-Fusion-Rv primer:

5’-TCAAGCTTGCTCTAGATTACAACTTGTTAAGCAAATC-3’;

CquiOR28-In-Fusion-Fw primer:

5’-AGATCAATTCCCCGGG<u>ACC</u>ATGGACGCCACCCAGCGA-3’ and

CquiOR28-In-Fusion-Rv primer:

5’-TCAAGCTTGCTCTAGATTACAGTTTGCTCAAAACGTC-3’.

Kozak sequence ACC (underlined) was added right before the start codon of the target gene. The colonies from transformation were verified by regular PCR with the cloning primers. The positive clones were cultured and subsequently extracted using the QIAprep Spin Miniprep kit (Qiagen) and sequenced by ABI 3730 automated DNA sequencer.

To identify introns for qPCR primer design, gDNA was extracted using Wizard Genomic DNA Purification Kit (Promega). Mosquitoes were homogenized individually with 600 μl of Nucleic Lysis Solution in ice-cold centrifuge tubes and placed at 65°C for 30 min. Subsequently, 3 μl of RNase was added, the mixture was incubated at 37°C for 30 min and finally cooled to room temperature for 5 min. Then, 200 μl of Protein Precipitation Solution was added, incubated on ice for 5 min, and centrifuged at 14,000 x g for 4 min. The supernatant was removed and transferred to a new tube containing 600 μl of isopropanol. After mixing, inverting, and centrifuging at 14,000 x g for 1 min, the supernatant was removed, and 600 μl of 70% ethanol was added. The gDNA pellet was carefully washed, the tube was centrifuged at 14,000 x g for 1 min, ethanol was aspirated, and the pellet was air-dried for 15 min. gDNA was reconstituted with 100 μl of DNA Rehydration Solution at 65°C, for 1 h. Lastly, gDNA was quantified by NanoDrop™ Lite Spectrometer (Thermo Fisher Scientific, Waltham, MA), and the quality was verified by agarose gel. The following primers were used to obtain the *CquiOR27* gDNA sequence.

gCqOR27Fw1: 5’-ATGGACACAGTCCGGTGGG-3’

gCqOR27Rv1: 5’-TTACAACTTGTTAAGCAAATCTTTGAGAACC-3’

gCqOR27Fw2: 5’-GGTTGCATATTCGAACAAATTTTGCTCACTATC-3’

gCqOR27Rv2: 5’-GATAGTGAGCAAAATTTGTTCGAATATGCAACC-3’

PCR products were extracted with Monarch PCR & DNA Gel Cleanup Kit (New England Biolabs, Ipswich, MA) and sequenced by ABI 3730 automated DNA sequencer.

### 2.3. qPCR

For quantitative PCR, RNA was extracted from antennal tissues with TRIzol reagent (Invitrogen), and cDNAs were synthesized from 200 ng of RNA of each sample using iScript™ gDNA Clear cDNA Synthesis Kit (Bio-Rad, Hercules, CA) according to the manufacturer’s protocol. Real-time quantitative PCR analyses were done by using SYBR™ Select Master Mix for CFX (appliedbiosystems by Thermo Fisher Scientific), a CFX96 Touch™ Real-Time PCR Detection System (Bio-Rad), and having *CquiRPS7* gene as a reference. The following primers were designed by the Primer 3 program (https://bioinfo.ut.ee/primer3-0.4.0/). Each forward (left) primer was manually selected to flank splicing junctions to make PCR products only from transcripts.

qCqRPS7-F: 5’-ATCCTGGAGCTGGAGATGA-3’

pCqRPS7-R: 5’-GATGACGATGGCCTTCTTGT-3’

qPCROR32-F: 5’-TCAAAGGGCTACGAATGGTC-3’

qPCROR32-R: 5’-GTGCGTCCAATACCGAAAGT-3’

qCqOR27-F: 5’-GATTTTTTCCGTGAAGTTGCTATAG-3’

qCqOR27-R: 5’-GCCAATGGCTGATCAAAACT-3’

qPCR was performed in three biological replicates, each having three technical replicates. Data were analyzed using the 2^-ΔΔCt^ method. Normalized data are displayed after dividing all 2-ΔΔCt values by the mean 2^-ΔΔCt^ for each group. One group was obtained with 5-day non-blood-fed mosquitoes and the other with 5-days after blood feeding, thus comparing relative expression levels of *CquiOR32* and *CquiOR27* in blood-seeking and oviposition-ready mosquitoes.

### 2.4. Oocytes preparations and two-electrode voltage-clamp recordings (TEVCs)

TEVC was performed as previously described (Leal et al., 2013; Pelletier et al., 2015; Xu et al., 2019; Xu et al., 2014; Xu et al., 2013; Xu et al., 2012a; Xu et al., 2012b; Xu and Leal, 2013; Xu et al., 2020; Xu et al., 2015; Zhu et al., 2013). In brief, linearized pGEMHE-CquiORs were used as templates to transcribe into capped cRNA with poly(A) using an mMESSAGE mMACHINE T7 kit (Ambion, Austin, TX) following the manufacturer’s protocol. The cRNAs were dissolved in RNase-free water and adjusted at a concentration of 200 μg/mL by UV spectrophotometry (NanoDrop™ Lite Spectrophotometer). 9.2 nl of a mixture of an OR and Orco cRNAs was microinjected into stage V or VI *Xenopus* oocytes (purchased from EcoCyte Bioscience, Austin, TX). Then, injected oocytes were incubated at 18°C for 3–7 days in modified Barth’s solution [in mM: 88 NaCl, 1 KCl, 2.4 NaHCO_3_, 0.82 MgSO_4_, 0.33 Ca(NO_3_)_2_, 0.41 CaCl_2_, 10 HEPES, pH 7.4] supplemented with 10 μg/mL of gentamycin, 10 μg/mL of streptomycin, and 1.8 mM sodium pyruvate. Test oocytes were placed in a perfusion chamber and challenged with a panel of odorants (see below). Currents were amplified with an OC-725C amplifier (Warner Instruments, Hamden, CT) holding the voltage at −80 mV and a low-pass filter at 50 Hz, and digitized at 1 kHz. Data acquisition and analysis were carried out with Digidata 1440A and pClamp10 software (Molecular Devices, LLC, Sunnyvale, CA).

### 2.4. Panel of odorants

The following compounds were used to challenge CquiOR27/CquiOrco- and CquiOR28/CquiOrco-expressing oocytes. Initially, the entire panel was screened with individual compounds at 1 mM diluted in perfusion buffer from their stock 1 M solutions. (-)-Caryophyllene oxide, (-)-menthone, (+)-limonene oxide, (+)-δ-cadinene, (±)-citronellal, (±)-lactic acid, (±)-linalool, (*E*)-2-heptenal, (*E*)-2-hexenal, (*E*)-2-hexenoic acid, (*E*)-2-hexenyl acetate, (*E*)-2-hexenyl acetate, (*E*)-2-methyl-2-butenal, (*E*)-2-nonenal, (*E*)-3-hexenoic acid, (*E*)-cinnamaldehyde, (*E,E*)-farnesol, (*E,E*)-farnesyl acetate, (*R*)-(+)-pulegone, (*S*)-(-)-perillaldehyde, (*Z*)-3-hexenyl acetate, (*Z*)-8-undecenal, 1,2-dimethoxybenzene, 1,4-diaminobutane, 1,5-diaminopentane, 1-butanol, 1-dodecanal, 1-dodecanol, 1-heptanol, 1-hepten-3-ol, 1-hexadecanol, 1-hexanol, 1-hexen-3-ol, 1-methylindole, 1-nonanol, 1-octanol, 1-octen-3-ol, 1-octen-3-one, 1-octen-3-yl acetate, 1-octyn-3-ol, 1-pentanol, 1-phenylethanol, 2,3-butanediol, 2,3-butanedione, 2,3-dimethylphenol, 2,4-dimethylphenol, 2,4-hexadienal, 2,4-thiazolinedione, 2,5-dimethylphenol, 2,6-dimethylphenol, 2-acetylthiophene, 2-butanol, 2-butanone, 2-butoxyethanol, 2-ethyltoluol, 2-heptanone, 2-hexanone, 2-hexen-1-ol, 2-methoxy-4-propylphenol, 2-methyl-2-thiazoline, 2-methyl-3-buten-2-ol, 2-methylindole, 2-methylphenol, 2-nonanone, 2-nonen-1-ol, 2-octanol, 2-octanone, 2-oxobutyric acid, 2-oxovaleric acid, 2-pentanol, 2-pentanone, 2-phenoxyethanol, 2-phenylethanol, 2-pyrrolidinone, 2-tridecanone, 2-undecanone, 3,4-dimethylphenol, 3,5-dimethylphenol, 3-hexen-1-ol, 3-hydroxy-2-butanone, 3-methyl-1-butanol, 3-methyl-2-butanol, 3-methyl-2-buten-1-ol, 3-methylbenzamide, 3-methylindole (skatole), 3-methylphenol, 3-octanol, 3-octyn-1-ol, 3-pentanol, 4,5-dimethylthiazole, 4-dimethylamino-1-naphthaldehyde, 4-ethylphenol, 4-methylcyclohexanol, 4-methylindole, 4-methylphenol, 5-hexanoic acid, 5-isobutyl-2,3-dimethylpyrazine, 5-methyl-2-hexanone, 5-methylindole, 6-methyl-5-hepten-2-one, 6-methylindole, 7-hydroxcitronellal, 7-methylindole, acetophenone, acetylacetone, benzaldehyde, benzyl alcohol, benzyl formate, butan-2-yl 2-(2-hydroxyethyl)piperidine-1-carboxylate (picaridin), butanal, butanoic acid, butyl acetate, butylamine, cadaverine, camphor, carbon disulfide, carvacrol, carvone, cinnamyl alcohol, citral, citronellol, cyclohexanone, cymene, decanal, decanoic acid, decyl acetate, dibutyl phthalate, dimethyl phthalate, dimethyl trisulfide, dodecanoic acid, ethanoic acid, ethyl 2-(*E*)-4-(*Z*)-decadienoate, ethyl 3-hydroxybutanoate, ethyl 3-hydroxyhexanoate, ethyl acetate, ethyl butanoate, ethyl hexanoate, ethyl lactate, ethyl linoleate, ethyl phenylacetate, ethyl propionate, ethyl stearate, ethyl *N*-acetyl-*N*-butyl-β-alaninate (IR3535), eucalyptol, eugenol, eugenyl acetate, farnesene, fenchone, furfural, geraniol, geranyl acetate, geranylacetone, guaiacol, heptanal, heptanoic acid, heptyl acetate, heptylamine, hexanal, hexanoic acid, hexyl acetate, hexylamine, indole, isobutyric acid, isopentyl acetate, isoprene, isopropyl myristate, isovaleraldehyde, isovaleric acid, jasmone, limonene, linalool oxide, linalyl acetate, linoleic acid, menthol, menthyl acetate, methyl (*N,N*)-dimethylanthranilate, methyl acetate, methyl anthranilate, methyl butyrate, methyl disulfide, methyl hexanoate, methyl myristate, methyl propionate, methyl salicylate, m-toluamide, myristic acid, *N*-(2-isopropylphenyl)-3-methylbenzamide, *N*-(sec-butyl)-2-methylbenzamide, *N,N*-diethyl-3-methylbenzamide (DEET), nerol, nerolidol, *N*-methylbenzamide, nonanal, nonanoic acid, nonyl acetate, *N*-sec-butyl-2-phenyl-acetamide, n-tridecanoic acid, ocimene, octadecyl acetate, octanal, octanoic acid, octyl acetate, octylamine, oleic acid, palmitic acid methyl ester, palmitoleic acid, p-coumaric acid, penatanal, pentanoic acid, pentyl acetate, pentylamine, phenethyl formate, phenethyl propionate, phenol, phenyl isobutyrate, phenyl propanoate, phenylacetaldehyde, phenylether, p-menthane-3,8-diol (PMD), propanal, propanoic acid, propyl acetate, propylamine, pyridine, pyrrolidine, sabinene, ß-caryophyphyllene, ß-myrcene, terpinen-4-ol, terpinolene, thujone, thymol, trimethylamine, trimethylamine, undecanal, α-hexylcinnamaldehyde, α-humulene, α-methylcinnamaldehyde, α-methylcinnamaldehyde, α-phellandrene, α-pinene, α-terpinene, α-terpineol, γ-decalactone, γ-dodecalactone, γ-hexalactone, γ-octalactone, γ-terpinene, and γ-valerolactone. The Orco ligand candidate 2-{[4-Ethyl-5-(4-pyridinyl)-4*H*-1,2,4-triazol-3-yl]sulfanyl}-*N*-(4-isopropylphenyl)acetamide (OLC 12 = VUAA-3) was used for confirming receptor protein expression.

### 2.5. Surface landing and feeding and hand-in cage assays

Repellency was measured using a surface landing and feeding assay described in detail elsewhere (Leal et al., 2017; Xu et al., 2014), with a slight modification. In short, this dual-choice assay had two Duddle bubbling tubes, which protrude from a wooden board inside of the arena (mosquito cage). Water was circulated inside the Duddle tubes to maintain their surface temperature at 37°C. A needle was placed on the top of each Duddle tube with the dual function of holding dental cotton rolls and delivering CO_2_ at 50 ml/min. Defibrinated sheep blood (100 μl) was applied to each cotton roll. To avoid contamination of the tubes with mosquito excretion, each tube was covered with a non-lubricated condom (Trojan ENZ) and replaced after five tests. Each Duddle tube was surrounded by a filter paper cylinder to which 200 μl of a test repellent solution or solvent only (control) was applied. Thus, one side of the arena (test) had a chemical curtain providing spatial repellency. Test mosquitoes (100 non-blood-fed, at least two weeks old) were deprived of water and sugar for at least one hour before the assays. DEET (1%) was used as a standard for comparison. Repellency tests were performed by testing DEET at one side of the arena, then DEET at the opposite side, two replicates of a test compound, one at each side of the arena, and continuing in this order for two or three complete cycles. Behavioral responses were expressed in protection rate according to WHO (WHO, 2009) and EPA (EPA, 2010) recommendations: P% = (C-T)/C) X100, where T and C represent the number of mosquitoes in the treatment and control sides of the arena after 5 min. The hand-in cage assay is a simplified version of the standard WHO (WHO, 2009) and EPA (EPA, 2010) arm-in cage assay. The four major differences are the cage’s size, hands vs. arm, the duration of the exposure, and starvation time: 1 h in vs. 12-24 h in WHO (WHO, 2009) and EPA (EPA, 2010) protocols. Collapsible field cages (30 × 30 × 30 cm; BioQuip Products, Rancho Cordova, CA) were used. One hundred to 200 non-blood-fed, 10-14 days old mosquitoes were used per cage. The test subject (W.S.L.) inserted a hand, which was covered with a glove, thus exposing only four fingers (see below). With a smaller cage, mosquitoes showed a robust response in a shorter time (1.5 min) than the duration recommended by WHO (WHO, 2009) and EPA (EPA, 2010), i.e., 3 and 5 min, respectively. Mosquitoes landing on the subject fingers for longer than 3 s were gently removed with a soft brush to prevent feeding. The UC Davis IRB was consulted and determined that IRB review was not required.

### 2.6. Formulation of γ-octalactone for the hand-in cage assay

A γ-octalactone cream formulation was prepared by adding stearyl alcohol (1.25 g) to 15 ml of water in a 50-ml beaker on a stirring hot plate and stirred with a magnetic bar until the alcohol melted. Next, 1.25 g of cetearyl alcohol was added and melted before adding γ-octalactone (1.75 g). Subsequently, polysorbate 80 (1.25 g), sorbitol oleate (1.25 g), stearic acid (1.25 g), glyceryl stearate self-emulsifying (1.25 g), and C12-C15 alkyl benzoate (0.5 g) were added in tandem after waiting to dissolve or melt. After 5 min, the mixture was cooled to 50°C before adding 0.25 ml of cyclomethicone. The resulting mixture was then transferred to a 50-ml conical centrifugal tube and let cool down to room temperature for subsequent use. For control, a similar cream formulation was prepared without γ-octalactone. The γ-octalactone cream formulation (ca. 1 g) was applied to the right hand’s small, ring, middle, and index fingers. The same amount of the control cream formulation was similarly applied to the left hand. Both hands were covered with Nitri-Solve 730-20 gloves, which were prepared by cutting the finger covers (except the thumbs), leaving only ca. 2 cm covering the proximal phalanx. Thus, the treated area had ca. 0.4 mg of γ-octalactone/cm^2^.

### 2.4. Statistical analysis and graphical preparations

Prism 9.3.1 from GraphPad Software (La Joya, CA) was used for both statistical analysis and graphical preparations. A dataset that passed the Shapiro-Wilk normality test was analyzed by t-test; otherwise, data were analyzed by Wilcoxon matched-pairs signed-rank or Mann-Whitney test, as specified below. All data are presented as mean ± SEM.

## 3. Results and discussion

### 3.1. CquiOR32/CquiOR27 relative expression

We cloned the full-length cDNA of CquiOR27 (GenBank OM240661) using primers designed based on a previously reported DNA sequence (Taparia et al., 2017). To design qPCR primers flanking splicing junctions, we cloned CquiOR27 gDNA (GenBank OM240662), given that the predicted RNA/mRNA sequence was not found in VectorBase. We re-analyzed our transcriptome data (Leal et al., 2013) and observed that TCONS_00034458 (Table S02) (Leal et al., 2013) is 97% identical to a segment (61%) of CquiOR27 DNA, with a 1% gap. We then compared by qPCR the relative expression levels of CquiOR32 and CquiOR27 at two different physiological stages, i.e., 5 days-old, non-blood-fed (host-seeking mosquitoes) and 5 days post blood feeding (oviposition-ready mosquitoes). qPCR analyses of *CquiOR28* were not performed because the receptor did not respond to any test compounds (see below). Although the transcription levels of the previously reported “inhibitory receptor” *CquiOR32* were higher than that of *CquiOR27* in both physiological stages (Fig. 1), *CquiOR27* showed relatively higher expression levels in non-blood-fed mosquitoes consistent with a possible role in host finding.

**Fig. 1.**
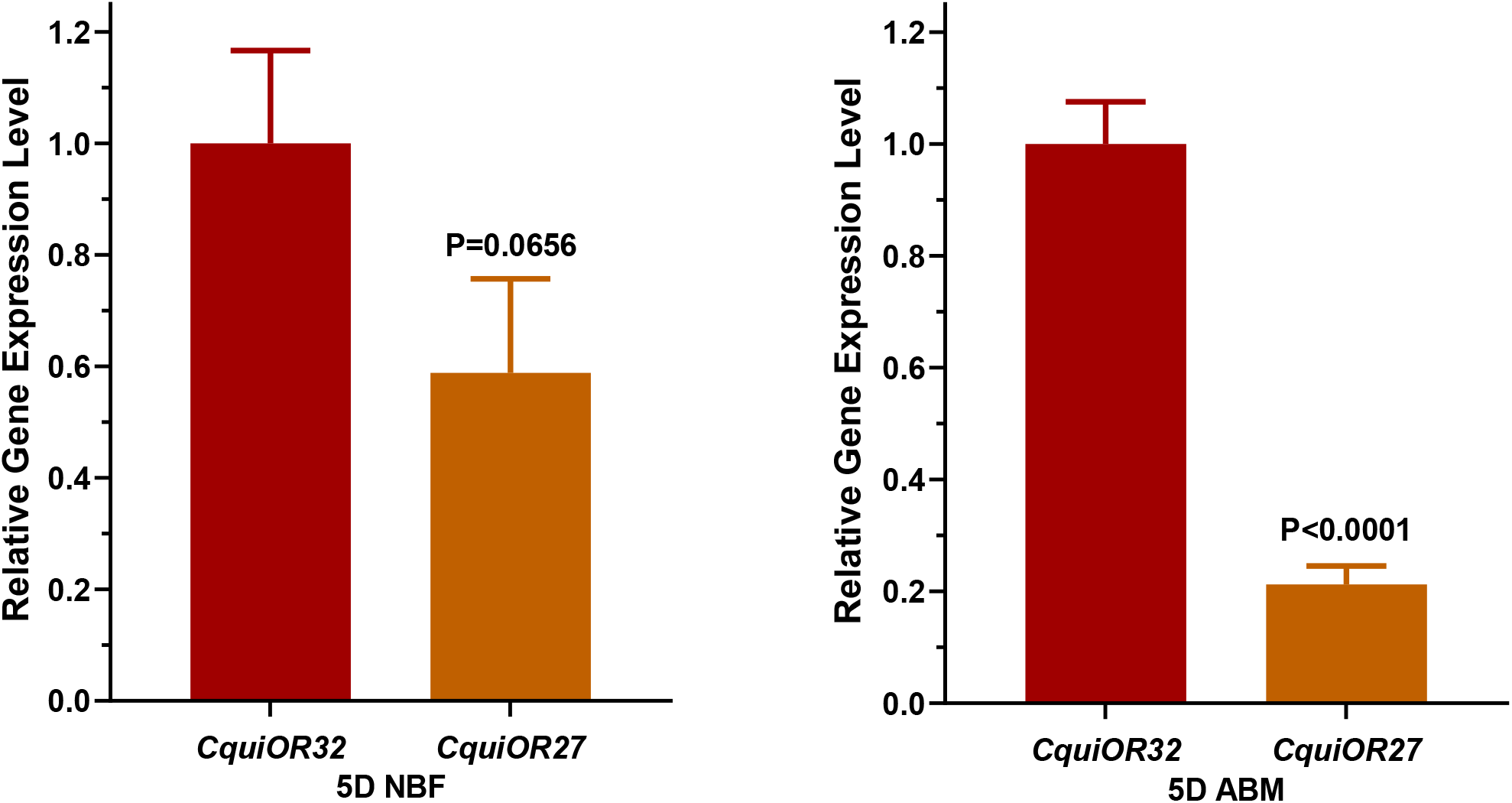
Relative transcript levels of *CquiOR32* and *CquiOR27*. Relative gene expression levels of *CquiOR*32 and *CquiOR27* in host-seeking and oviposition-ready mosquitoes. 5D, 5 days; NBF, non-blood-fed; ABM, after a blood meal.

### 3.2. CquiOR27 generated regular and inhibitory currents

To deorphanize CquiOR27, we used a panel of 250 odorants and two-electrode voltage-clamp recordings from *Xenopus laevis* oocytes co-expressing CquiOR27 and CquiOrco. Small and moderate inward currents were elicited by fenchone, citronellal, alpha-terpineol, *N*-sec-butyl-2-phenyl-acetamide, camphor, ethyl linoleate, 1-phenylethanol, 2,3-dimethylphenol, 3,5-dimethylphenol, jasmone, octadecyl acetate, limonene oxide, carvone, (*R*)-(+)-pulegone, and menthyl acetate. Additionally, three compounds, linalool oxide, γ-octalactone, and γ-decalactone, elicited robust responses (Fig. 2). By contrast, 2,6-dimethylphenol, 3,4-dimethylphenol, 2,4-dimethylphenol, 2,3-dimethylphenol generated inhibitory currents, although the strongest responses were elicited by octylamine (Fig. 2). Of note, two phenolic compounds (2,3-dimethylphenol and 3,5-dimethylphenol) elicited inward currents, whereas others showed inhibitory responses.

**Fig. 2.**
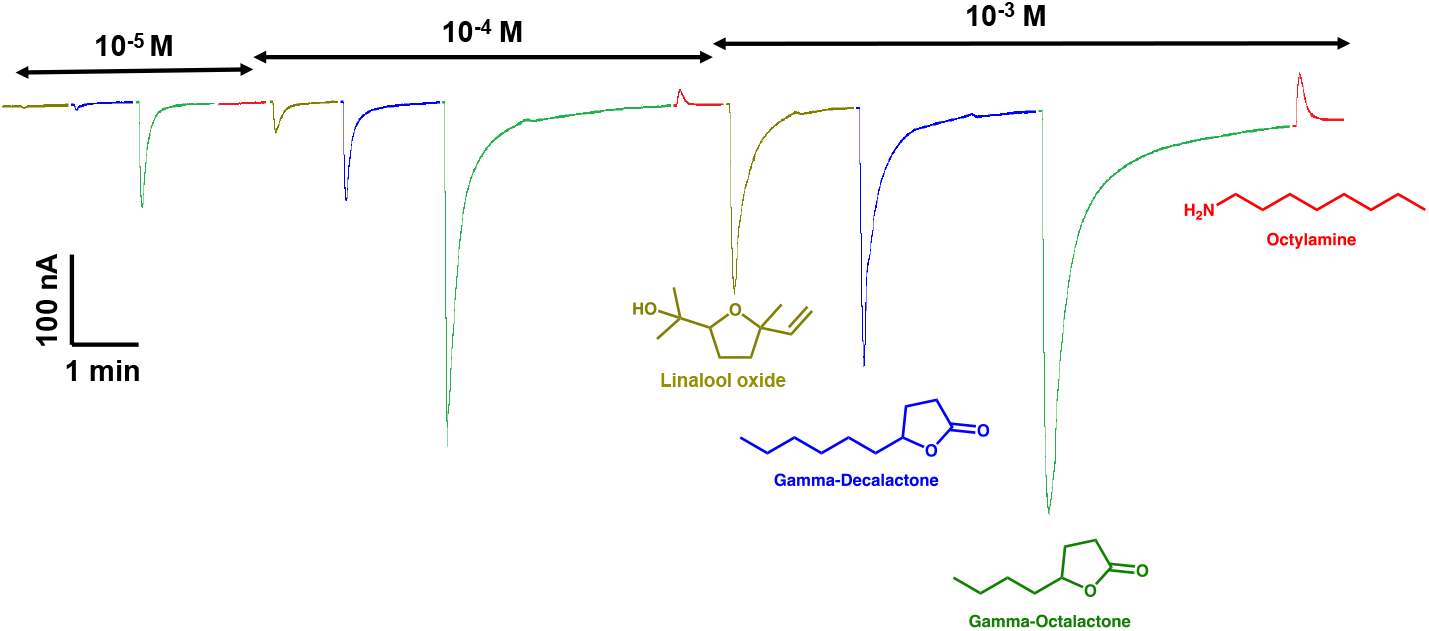
Currents recorded from CquiOR27/CquiOrco-expressing oocytes with stimulatory and inhibitory compounds. Representative trace of the responses elicited by three main stimulatory and one inhibitory compounds, i.e., linalool oxide, γ-octalactone, γ-decalactone, and octylamine. This trace was colored to match the colors of the corresponding structures. Dose increased from left to right, from 0.01 to 1 mM.

Although CquiOR27 showed a promiscuous profile, γ-octalactone and γ-decalactone were by far the best ligands in our screenings (Fig. 3), with a moderate response to linalool oxide. Interestingly, the responses to γ-hexalactone and γ-dodecalactone were very small. The dose-response relationship confirmed that indeed γ-octalactone was the best ligand (Fig. 4).

**Fig. 3.**
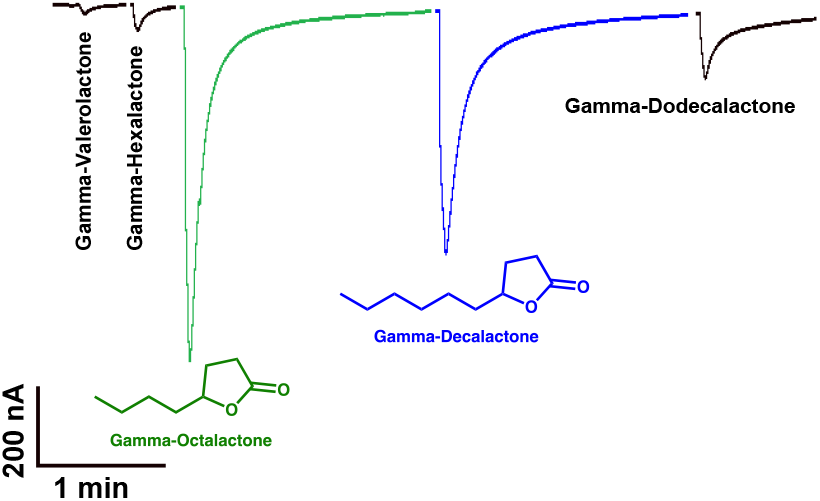
Currents recorded from CquiOR27/CquiOrco-expressing oocytes with lactones. Recordings obtained with five γ-lactones at the same dose (1 mM), with the same oocyte preparation.

**Fig. 4.**
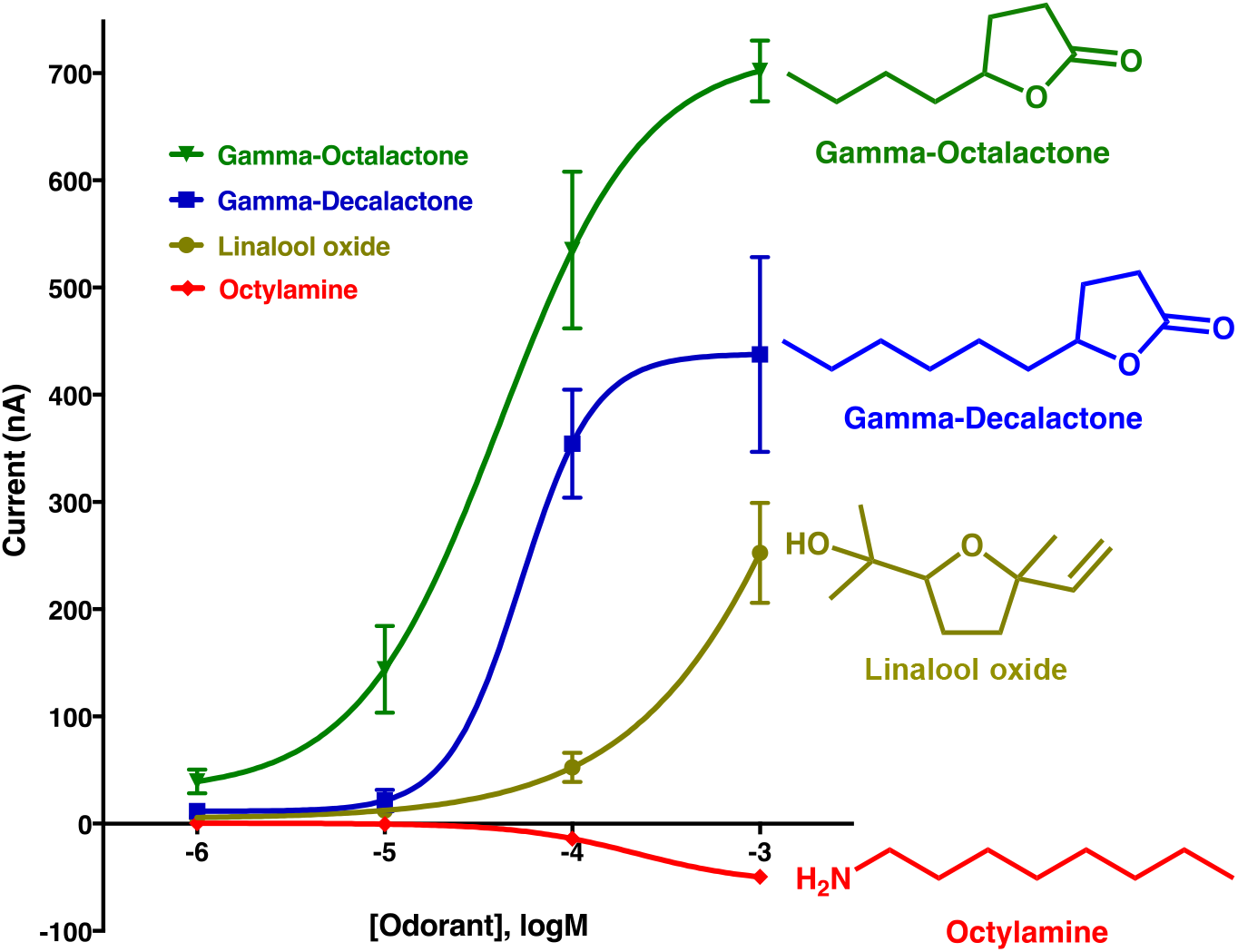
Concentration-dependent curves obtained with the main stimulatory and inhibitory compounds. Recordings were obtained with five CquiOR27/CquiOrco-expressing oocytes (N = 5). Each oocyte was challenged with all four compounds starting with a dose of 10^−6^ M (1 μM) and increasing to 10^−3^ M (1 mM). The best ligands were γ-octalactone (EC_50_ 4.1 × 10^−5^ M) and γ-decalactone (EC_50_ 5.3 × 10^−5^ M).

### 3.3. Intrarecptor inhibition manifested in the *Xenopus* oocytes recording system

We challenged CquiOR27/CquiOrco-expressing oocytes with excitatory and inhibitory compounds. Specifically, we presented γ-octalactone, octylamine, and a mixture of the two compounds. Octylamine per se generated strong inhibitory currents (upward deflections), whereas γ-octalactone alone gave robust stimulatory currents. The γ-octalactone-generated inward currents decreased significantly in a dose-dependent manner when this odorant was co-applied with octylamine (Fig. 5). Oocyte adaptation was ruled out as there was no significant difference between the γ-octalactone-elicited responses before and after co-applications with octylamine (p=0.9683, Mann-Whitney test). Similar findings were observed when 2,3-dimethylphenol was used as the inhibitory compound (Figs. 1). We, therefore, concluded that both stimulatory and inhibitory compounds are acting on the same CquiOR27/CquiOrco complex, thus suggesting an intrareceptor interaction.

**Fig. 5.**
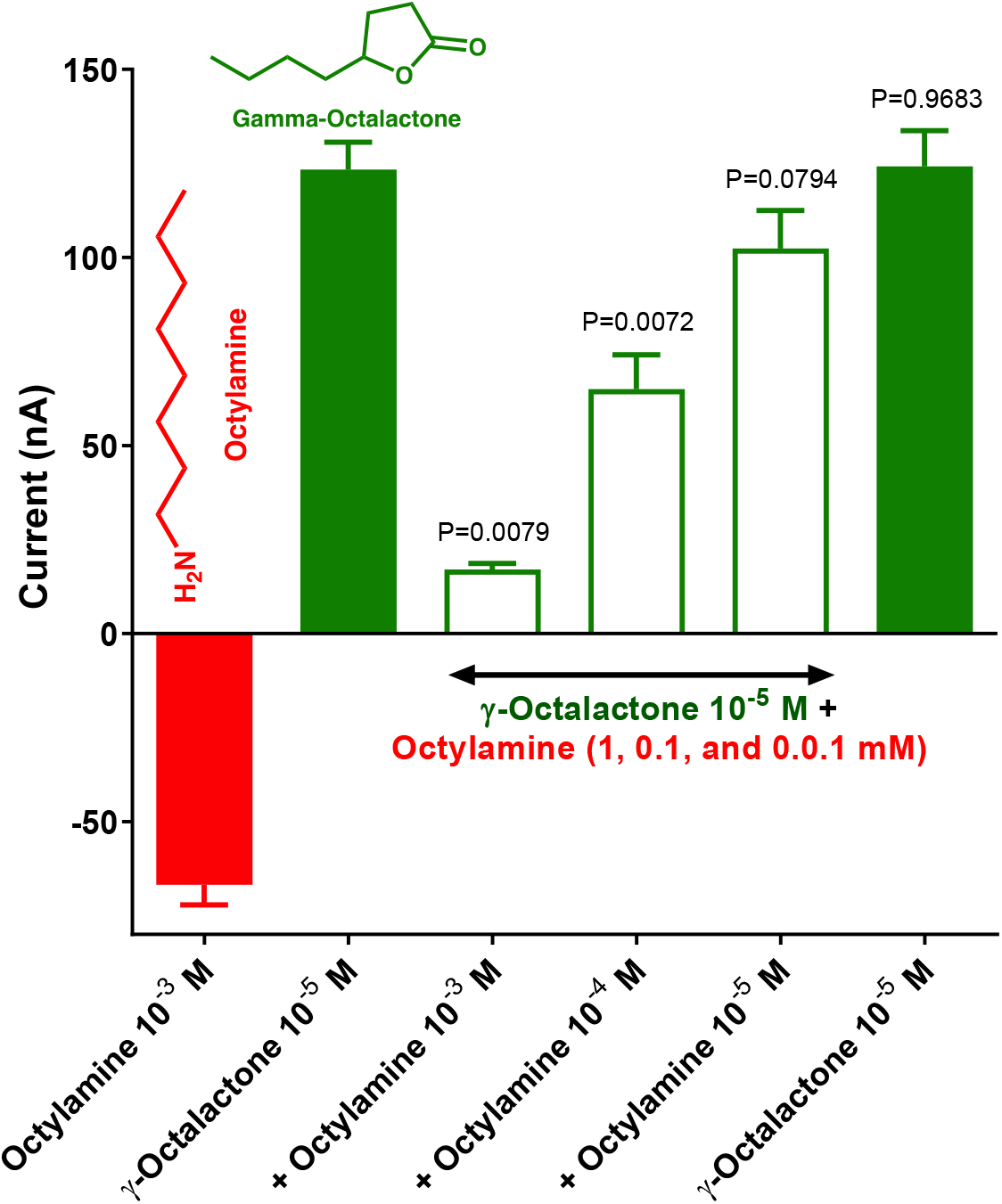
Quantification of responses to binary mixtures of stimulatory and inhibitory compounds. Quantification of responses recorded from CquiOR27/CquiOrco-expressing oocytes when challenged with octylamine and γ-octalactone separately or combined. Each oocyte was challenged with the inhibitory compound at 1 mM, γ-octalactone at 0.1 μM and then a mixture of γ-octalactone at 0.1 μM and the inhibitory compound in decreasing doses from 1 mM to 0.01 mM and finally γ-octalactone at 0.1 μM. Recordings were obtained with 5 CquiOR27/CquiOrco-expressing oocytes.

As previously discussed (Xu et al., 2019), there are at least two possible explanations for the currents in the reverse direction elicited by octylamine and other inhibitory compounds. These ligands may be negative allosteric inhibitors, or perhaps they are inverse agonists. It is conceivable that by binding to CquiOr27/CquiOrco-complex, the open <=> close receptor equilibrium shifts towards the close, inactive conformation, thus suppressing signaling. In the case of CquiOR32 (Xu et al., 2019), we reasoned that, given the different structures, stimulatory and inhibitory compounds might be acting on different binding sites. This argument is still valid when comparing γ-octalactone with octylamine. However, the observation that isomers of dimethylphenol gave inward and outward currents suggests that the compounds eliciting currents in the reverse direction might be inverse agonists.

### 3.4. CquiOR28 is non-functional

We selected and sequenced 32 colonies of CquiOR28. They gave four variants (OM258698-OM258701), which were subcloned in pGEMHE and, subsequently, tested in the *Xenopus* oocyte recording system. No stimulatory or inhibitory currents were recorded when these variants were challenged with our panel of odorants. Because none of the variants responded to Orco ligand candidates (VUAA-1 or OCL12), we concluded that CquiOR28 is not a functional receptor.

### 3.5. γ-Octalactone is a potent repellent

Given that γ-dodecalactone and related compounds have been previously reported as mosquito repellents (Bedoukian, 2016), we tested the repellency activity of the major ligands for CquiOR27 against *Cx. quinquefasciatus*. Specifically, we measured γ-octalactone and γ-decalactone repellency against *Cx. quinquefasciatus* using the surface landing and feeding assay (Leal et al., 2017; Xu et al., 2014) and compared their potency with the gold standard of insect repellents, DEET. Repellency activity was expressed in protection, according to WHO (WHO, 2009) and EPA (EPA, 2010) recommendations. Of note, protection (P) differs slightly from the repellency index (RI) commonly used in insect behavior. These indexes are obtained by dividing the responses to (control minus treatment) by the responses to control only (WHO and EPA protocol) or (control plus treatment) in the generic repellency index. Specifically, P=(C-T)/C and RI=(C-T)/(C+T), respectively. Thus, when T is very small, as in this case, the two indexes are nearly equal.

The protection elicited by 1% γ-octalactone (91.2 ± 2.3%) was not significantly different from the protection obtained with the same dose of DEET (90.0 ± 2.8%) p=0.2416, Wilcoxon matched-pairs signed-rank test). Similarly, γ-decalactone (85.9 ± 3.5 %) did not differ significantly from DEET (85.0 ± 2.1 %, p=0.8438). We then directly compared γ-octalactone with γ-decalactone by measuring their repellency with the same test mosquitoes. The protection elicited by γ-octalactone (89.0 ± 2.6 %) was not significantly different from that measured with γ-decalactone (89.3 ± 2.6 %, N=25, p=0.9935, Figs 2). In short, γ-octalactone did not differ significantly from γ-decalactone, and these two natural repellents did not differ from DEET in our assays. By contrast, γ-dodecalactone (65.7 ± 6.1 %) showed a significantly lower repellency activity than DEET (85.8 ± 4.6 %, N=10, p=0.0171, paired, two-tailed t-test).

Previously, delta-octalactone has been reported as a repellent against the tsetse fly, *Glossina morsitans* (Mwangi et al., 2008). Still, a literature survey (PubMed and Google Scholar) did not yield any evidence that γ-octalactone has been previously reported as a mosquito repellent. This is surprising because of the extensive prospect of repellent from natural sources (Gross and Coats, 2014; Paluch et al., 2010).

Given the strong responses observed in the surface landing and feeding assay, we next tested γ-octalactone in a modified hand-in-cage assay. We designed a simplified version of the arm-in-cage assay recommended by WHO (WHO, 2009) and EPA (EPA, 2010). With a smaller cage, mosquitoes readily landed on the untreated subject’s fingers, thus allowing a shorter (1.5 min) testing time. For these hand-in cage assays, we used a 7% cream formulation, consistent with the lowest percentage of DEET products in the market. Mosquito responses to one hand of the subject treated with a γ-octalactone cream formulation were compared with the responses to the other hand covered with a control cream devoid of γ-octalactone (untreated). γ-Octalactone provided 97.0 ± 1.3% protection (Fig. 6), with 47.6 ± 8.3% and 1.3 ± .7% landings per trial in the untreated and treated hands, respectively (N=8, p=0.0078, Wilcoxon matched-pairs signed-rank test).

**Fig. 6.**
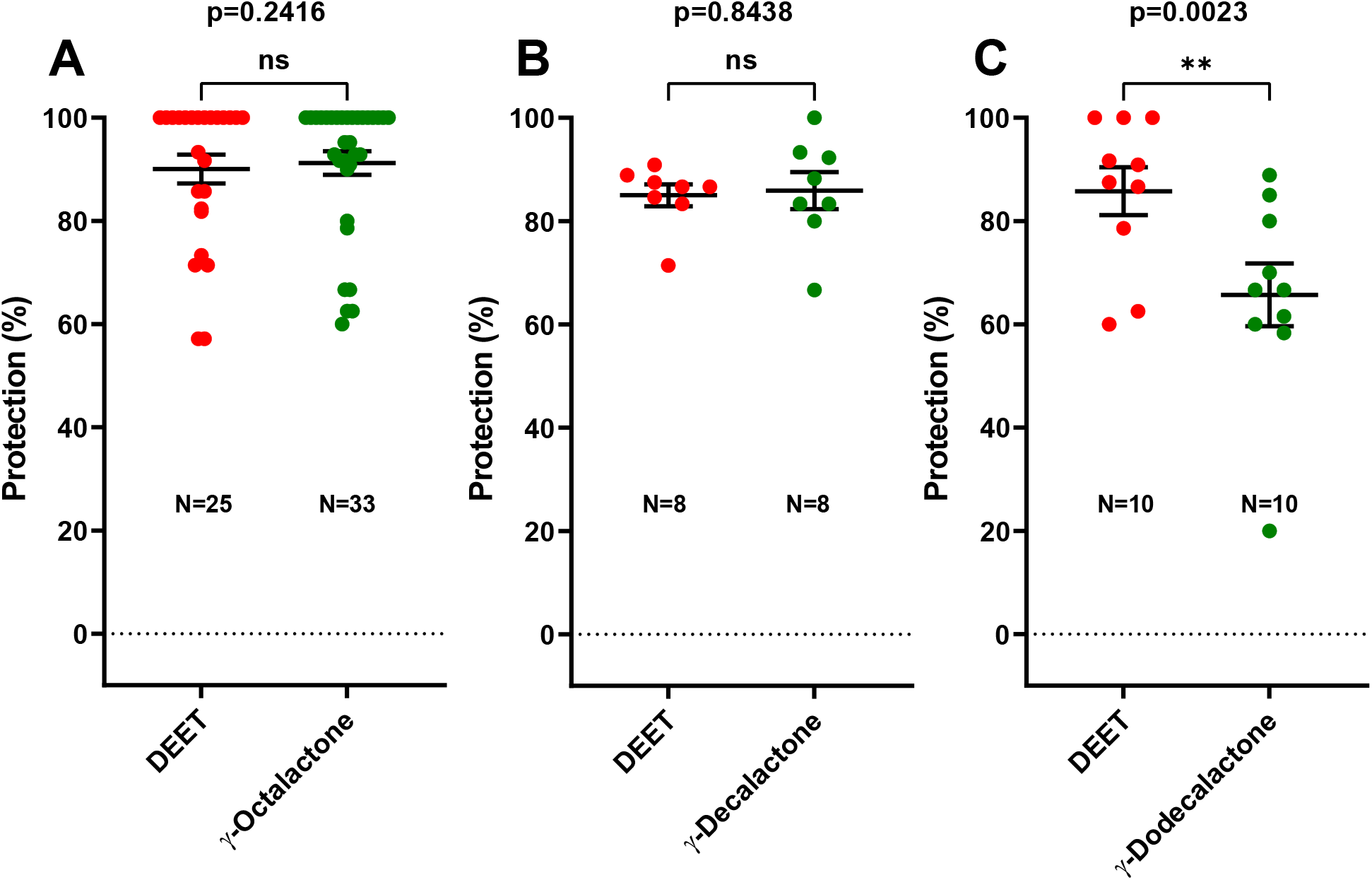
Behavioral responses of *Cx. quinquefasciatus* mosquitoes in repellency assays. Comparative responses of non-blood-fed female mosquitoes to DEET, γ-octalactone, γ-decalactone, and γ-dodecalactone in the surface landing and feeding assay. Repellency activities are expressed in the protection rate. The repellency activities of γ-octalactone (A) and γ-decalactone (B) did not differ significantly from that of DEET, but γ-dodecalactone showed significantly lower protection than DEET (C).

**Fig. 7.**
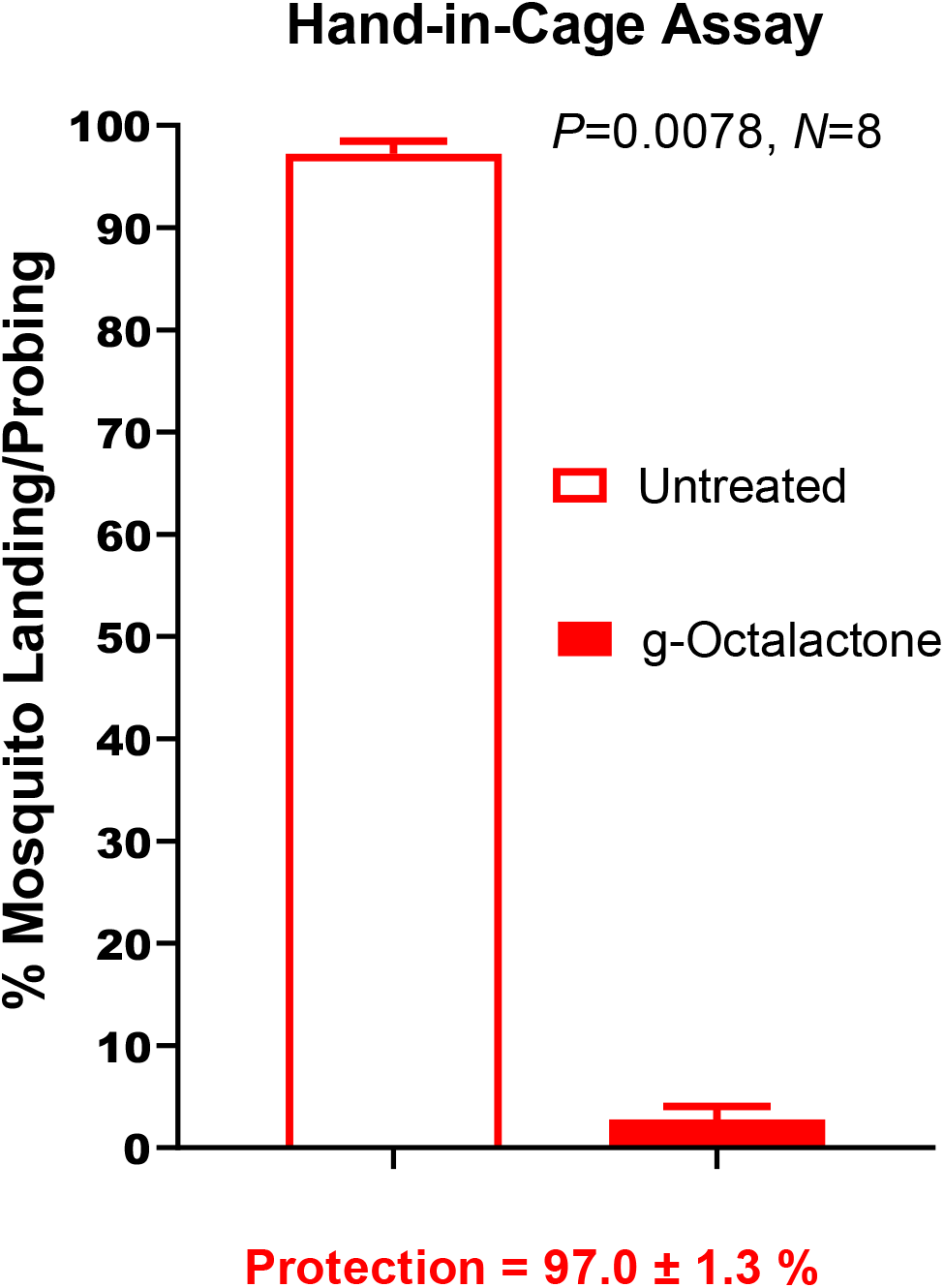
γ-Octalactone repellency measured in a hand-in cage assay. Responses of mosquitoes (landing/probing) to hands covered with a γ-octalactone cream formulation and a control formulation devoid of γ-octalactone (untreated).

### 3.4. Overall conclusions

Our reverse chemical ecology approach led to discovering a potent spatial natural repellent of potential practical applications. We showed that γ-octalactone is a strong repellent, but it is necessary to develop a slow-release formulation for complete long-term protection. Of note, γ-octalactone is a natural compound found in peaches, mangoes, beef, and ham (PubChem) used in hand creams and as a flavoring agent.

We found evidence in these studies that, like CquiOR32 (Xu et al., 2019), CquiOr27 is a receptor with dual current directions (“bidirectional”). Specifically, CquiOR27/CquiOrco-expressing oocytes were activated by some compounds, including γ-octalactone, and inhibited by other ligands, including octylamine. It is conceivable that the observed currents in the reverse direction, elicited by octylamine and some phenolic compounds, are not outward currents sensu stricto. The observed currents in the reverse direction could be derived from “baseline shift” due to the closing of ion channels (reduction of spontaneous activity) upon CquiOR27 binding to inhibitory compounds. To be rigorously tested, these hypotheses must await the elucidation of the 3D structures of these receptors.

## Author contributions

P.X., Y-M. C., S. A., G.M.L. and W.S.L. performed research; W.S.L. designed research; P.X. and WS.L. analyzed the data; W.S.L. wrote the paper; all authors read and approved the final manuscript.

## Acknowledgments

We thank Erica A. Littman for the preliminary γ-octalactone repellency tests. The Orco ligand candidate OCL12 was provided by the Vanderbilt Institute of Chemical Biology, Chemical Synthesis Core, Vanderbilt University, Nashville, TN 37232-0412.

## Supplementary information

Supplementary figures: Figs1 and Figs2

## Supplementary figure legends

Figs1. **Quantification of responses to binary mixtures of stimulatory and inhibitory compounds**

Quantification of responses recorded from CquiOR27/CquiOrco-expressing oocytes when challenged with 2,3-dimethylphenol and γ-octalactone separately or combined. Each oocyte was challenged with the inhibitory compound at 1 mM, γ-octalactone at 0.1 μM and then a mixture of γ-octalactone at 0.1 μM and the inhibitory compound in doses from 1 mM to 0.01 mM and finally γ-octalactone at 0.1 μM. Recordings were obtained with 3 CquiOR27/CquiOrco-expressing oocytes.

Figs2. **Direct comparison of protection provided by γ-octalactone and γ-decalactone**

There was no significant difference between the repellence activities of γ-octalactone and γ-decalactone as measured by direct comparison in the surface landing and feeding assay (N = 25, P=0.9935, Wilcoxon, two-tailed, matched-pairs signed-rank test.

