## Supplementary figures and images for "Mosquito odorant receptor sensitive to natural spatial repellents and inhibitory compounds"

### Supplemental Figures 1 & 2

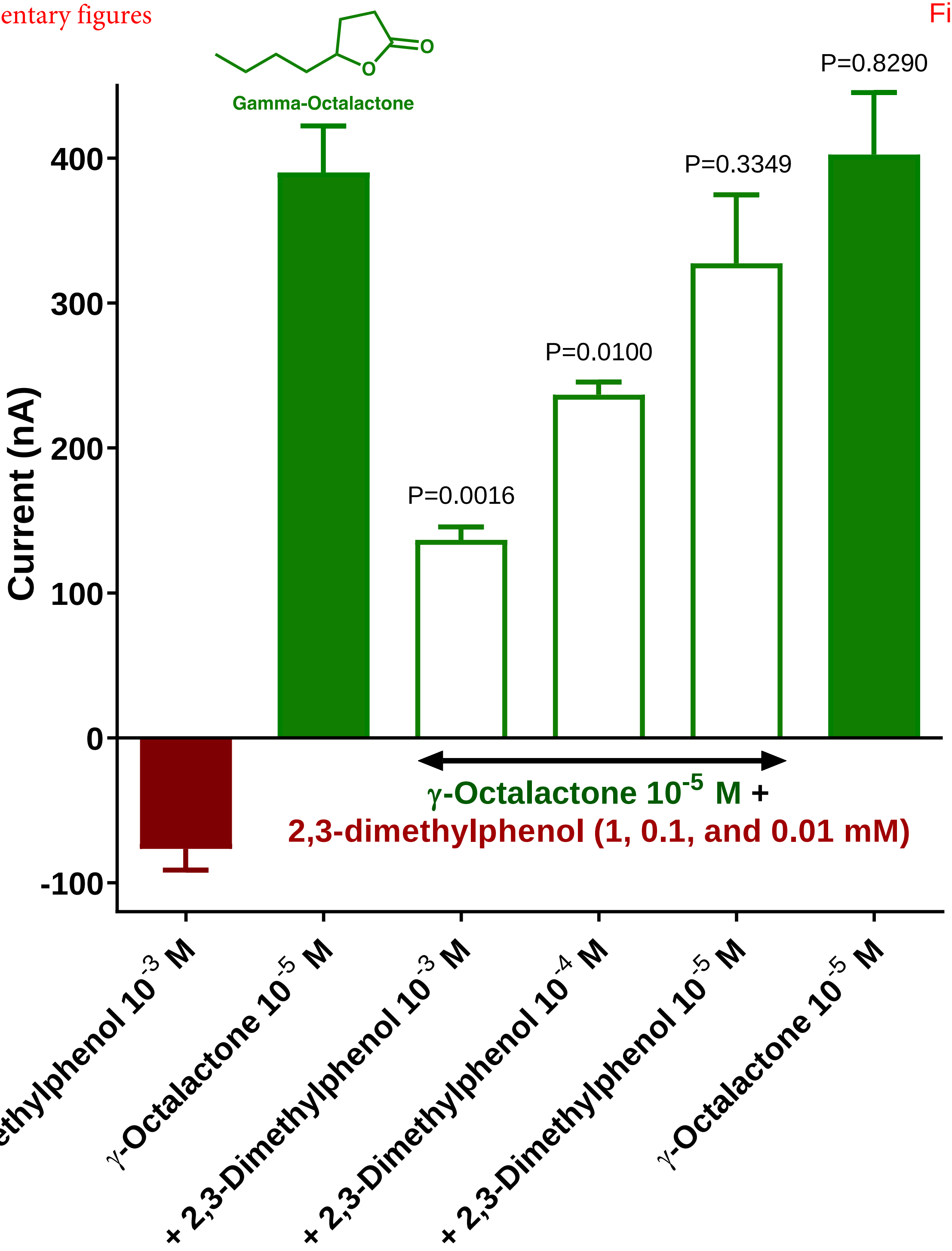

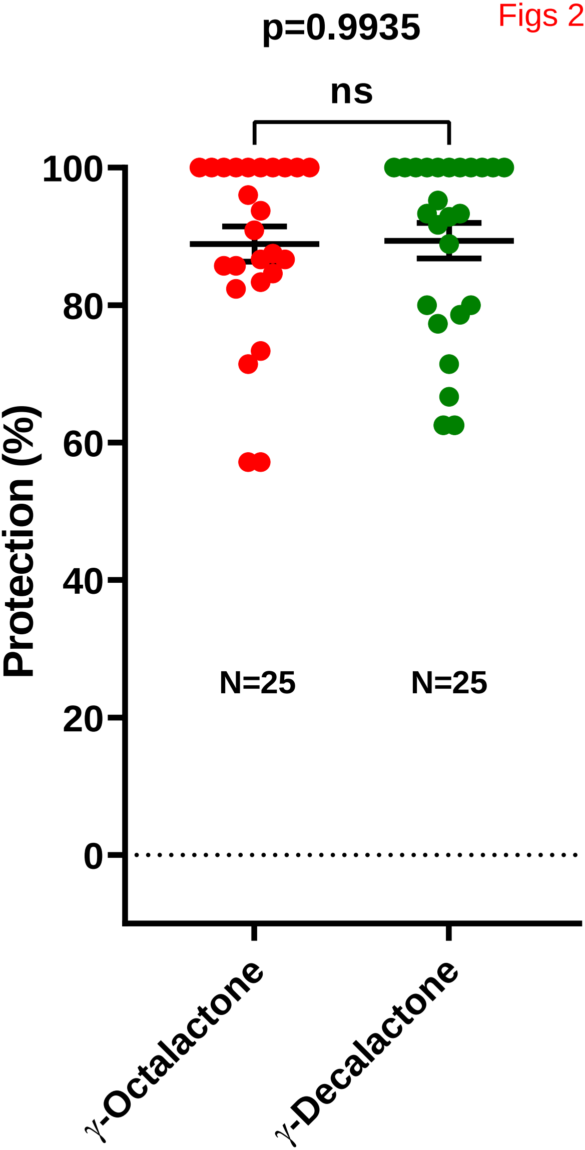
